# The Novel Chaperone Activity of p47 Modulates Force-Mediated Retrotranslocation from the Endoplasmic Reticulum to the Cytosol

**DOI:** 10.1101/2024.12.30.630725

**Authors:** Deep Chaudhuri, Shubhasis Haldar

## Abstract

Protein quality control in the endoplasmic reticulum (ER) maintains cellular proteostasis by regulating protein folding, assembly, and degradation. While protein translocation from the cytosol into the ER is well-studied, the reverse process, known as retrotranslocation, which exports proteins from the ER lumen or membrane into the cytosol, is less understood.

In this study, we investigate the role of p47, a cofactor of the ATPase p97, in retrotranslocation and its chaperone-like activity on the model substrate talin using single-molecule magnetic tweezers. We find that p47 enhances the mechanical stability of talin, by increasing its folding probabilities by 8 pN, which generate a maximum work output of up to ∼199.5 zJ to facilitate the extraction of the protein from the ER lumen. These findings suggest that p47 interacts with substrates emerging from the ER, generating pulling forces that aid retrotranslocation, which uncovers a novel mechanistic role for p47 in ER-associated degradation, advancing the understanding of retrotranslocation.

## Introduction

Protein quality control in the endoplasmic reticulum (ER) is an essential process for maintaining cellular homeostasis by ensuring proper folding, assembly and degradation of proteins^1–4^. This process relies on a well-coordinated mechanism of forward translocation and retrotranslocation^5,6^. In forward translocation, protein can be translocated across the membrane by pulling or sterically directed their movement^5,7^. In this aspect, molecular chaperone such as Bip facilitate the unidirectional movement of the unfolded substrate on the lumen side of the membrane^8^. However, Bip is unlikely to function in reverse during retrotranslocation. Retrotranslocation, a critical step in ER associated degradation (ERAD) pathway, extracts the misfolded proteins from the ER and delivers them to the cytosol for proteasomal degradation. Several studies have implicated the AAA family of ATPases, particularly Cdc48 in yeast and its mammalian ortholog p97, in driving the extraction of polypeptides from the ER membrane^9–12^. The precise mechanisms underlying the extraction of the unfolded substrate and the potential involvement of additional chaperones or cofactors facilitating this force-driven retrotranslocation remain poorly understood. Mechanical force recognized as critical regulators of protein translocation processes, particularly in post-translational pathways. For instance, in mammalian cell, Sec 61 tunnel associated chaperone Bip and ERdj assist the translocating of the unfolded substrate by promoting substrate folding at higher force range and preventing the backsliding of the polypeptide during the translocation through Sec61 translocon pore^13^. In bacterial systems, the ATPase SecA generates ∼10 pN of mechanical force to translocate the substrate protein through the Sec YEG tunnel, while co translational folding at the ribosomal exit tunnel occurs under mechanical tension, facilitating secretory protein transport^14^. These studies highlight the importance of mechanical force for pulling the unfolded substrate from tunnel during the protein translocation process. In this context, retrotranslocation involves the extraction of the unfolded substrate from ER to cytosol, a reverse directional process raises the intriguing question of whether mechanical forces play similar role in driving retrotranslocation.

Here we have tries to understand the independent roles of p47, a known cofactor of p97, as a potential regulator during retrotranslocation and its chaperone activity in this process. Previous studies have demonstrated that p47 modulates the ATPase activity of p97 and bind more strongly to the ATP bound form of p97, suggesting it may have a role in force generation in the retrotranslocation process^12,15^. Our custom made advanced single molecule force spectroscopic technique, magnetic tweezers enable precise measurement of the folding unfolding dynamics of a single protein molecule across extensive force range of 4-120 pN, with exceptional special and temporal resolution^16–19^. Using our approach, we can be able to apply the force on the protein substrate through both force ramp and force clamp protocols. The force ramp protocols enable us to investigate the mechanical strength and the kinetic parameters of protein molecules at varying pulling velocities with minimizing thermal fluctuations^19^. In contrast, the force clamp technique allows us to explore the folding-unfolding dynamics, conformational change and polymeric properties of the protein molecules^16,17^. The advantage of long force range allows us to investigate the properties of a single protein at sub-pN resolution.

Using this single-molecule techniques, we tried to understand the independent role of p47 in retrotranslocation, shedding light on its mechanism for extracting substrates from the ER to the cytosol. Here we used talin, a well-characterized mechanosensitive model protein with well-defined two-state folding dynamics under force, an ideal noble substrate, enabling us to precisely measure the effects of p47^19–22^. Our results demonstrated that p47 significantly enhances the folding probability of the talin at higher force regions, shifting the half point force from 8.3 pN to 16.6 pN. We further investigated the mechanical strength of talin by quantifying its average unfolding and refolding force. This finding reveals that the presence of p47 enhances the mechanical stability of the substrate, showing a consistent increase in stability across constant pulling velocities. Additionally, our analysis demonstrated that p47 enhances the maximum mechanical work out put, reaching approximately 199.5 zJ of mechanical work under mechanical stress, facilitating efficient extraction of the polypeptides from ER to the cytosol. This finding suggests a mechanistic model in which p47 associated to the ER translocation machinery, interacts with the substrate emerging from the ER lumen and generates additional pulling force to facilitate retrotranslocation from ER to the cytosol. Overall, our findings highlight a potential connection between p47’s chaperone-like activity and its role in the ERAD pathway, advancing our understanding of the molecular machinery that safeguards cellular proteostasis.

## Results

### Mechanical stability and folding Dynamics of the talin R3 IVVI domain observed using single-molecule magnetic tweezers

The mechanical stability and the conformational transition of the talin R3 IVVI domain has been investigated at single molecule resolution using our custom build covalent magnetic tweezers. The application of this technique allows us to understand the folding-unfolding dynamics of a single protein under sub-pN force resolution. Fig. 1A depicts the representative experimental set up for magnetic tweezers experiment, where C-terminal end of the protein molecule is attached with the Avi tag for the precise biotin streptavidin attachment with the magnetic bead and N-terminus of the protein molecule is attached with the glass surface using Halo tag chemistry^16,17,23,24^. A permanent magnet is positioned just above the glass chamber and the force is applied on the paramagnetic bead attached with the substrate protein. By adjusting the distance between the permanent magnet and the magnet attached to the glass surface, the applied force on the protein sample can be precisely controlled. The details of the force calibration method have been discussed in our previous study^16,17,24,25^.

**Figure 1:**
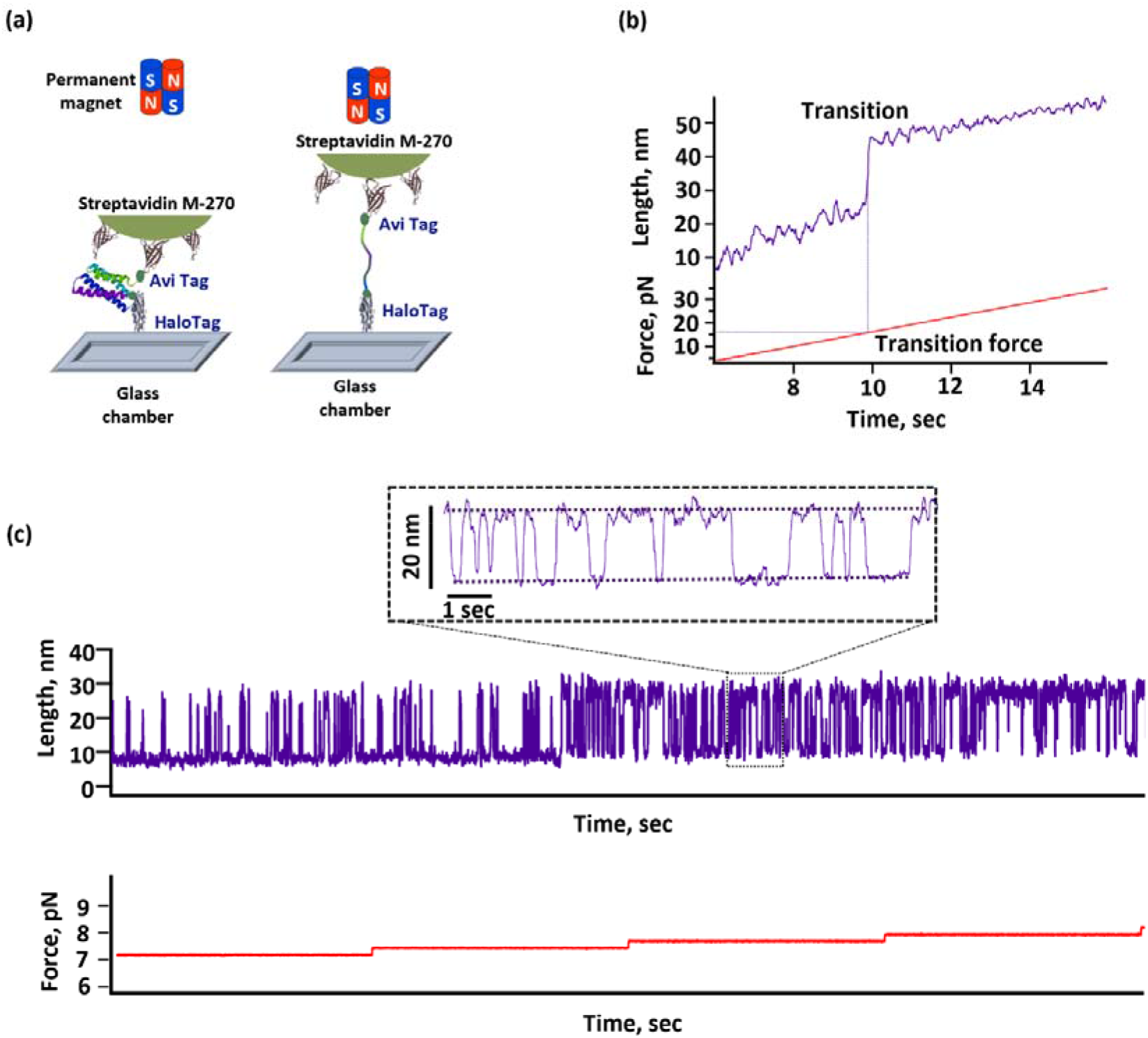
Representative experiment set up and trace obtained for magnetic tweezers experiment: **(A)** Schematic representation of the magnetic tweezers experiments. The C terminus of the protein is tethered to a streptavidin coated magnetic bead and N terminus of the protein is attached to the glass surface via HaloTag chemistry. The applied force can be controlled by adjusting the distance in between the two magnets. **(B)** Representative trace for force ramp protocol, where the force has been increased from 4 to 35 pN at a constant velocity of 3pN/sec. The unfolding transition force was determined by extrapolating the transition step to the force ramp axis. **(C)** Representative trace for the foding dynamics of talin R3 IVVI domain. This trace characterised the folding unfolding population of talin R3 IVVI domain, revealing the populations of folded and unfolded states at varying forces. The inset highlights the equilibrium state of talin at a force of 8 pN, where the protein is equally distributed in the folded and unfolded state.

This single molecule technology allows two distinct experimental approach force ramp and force clamp technology. Fig. 1B illustrate the application of force ramp technique to understand the mechanical stability of the protein molecule. Here force is applied incrementally at a particular rate of 3 pN/sec to observe the transition force of the substrate protein where the protein jumps from folded state to the unfolded state, reflects the mechanical strength of the protein molecule. The force-dependent transition behaviour of talin can be modulated through interactions with different proteins or chaperone molecules, which may influence the unfolding and refolding forces observed using force-ramp protocols. Force clamp technique provides us to apply a constant force on the substrate molecule, enable us to understand the transition in between the folded and the unfolded state. Fig. 1C displays a complete representative trajectory of R3 IVVI folding dynamics, showing the population of both folded and unfolded state at applied unfolding forces. The folding dynamics was determined by analysing the relative population of the folded state of multiple folding trajectories based on the total duration of the observed transitions. At 7 pN, talin R3 IVVI domain is predominantly in the folded state. However, upon increasing the force up to 8 pN, the protein hops in between the folded and unfolded state, exhibiting a transition step of ∼20 nm.

### p47 mediates change on the folding probability talin R3 IVVI domain

Fig. 2 illustrates the change in the folding probability of talin in the presence of p47. Interestingly, our findings revealed that the folding probability of talin has been shifted towards the higher force regions in the presence of 10 µM p47. Here we have found that the half point force (the force at which the protein is equally populated both in the folded and unfolded state) of talin was increased significantly from 8.61 pN to 16.6 pN in the presence of p47. We have further calculated the half point force for talin at varying concentrations of p47 and observed that with increasing the concentration of p47 the half point force of talin is increased and it’s become saturated after 10 µM (Supp. Fig. 1).

**Figure 2:**
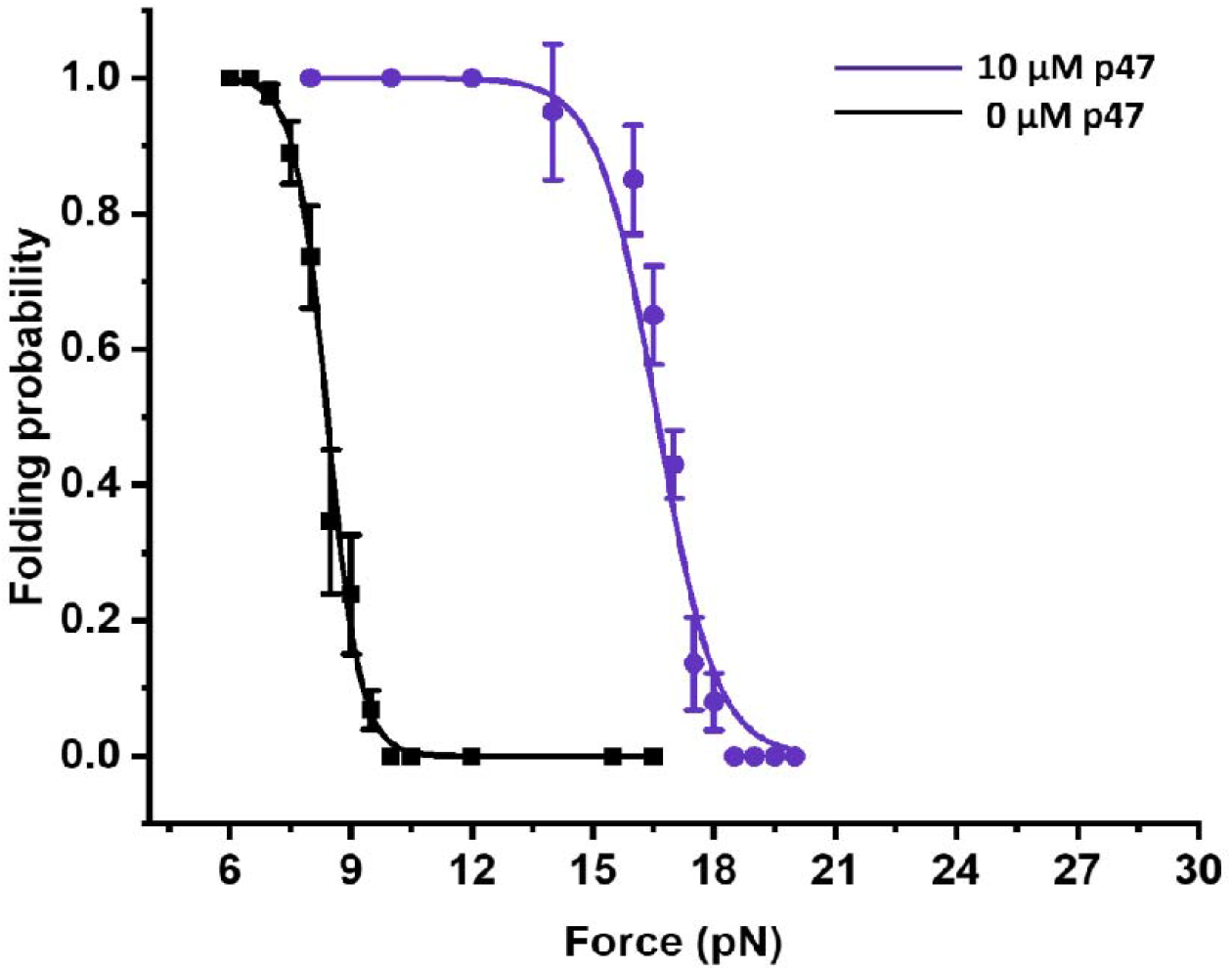
Change in the folding probability of talin in the presence of p47: Folding probability of talin has been plotted as a function of force. It has been observed that the half point force, where talin is equally populated in folded and unfolded stat, was determined to be 8.4 pN (black). In the presence of p47 (purple), this value significantly shifted to 16.6 pN. Each data points represented measurement from >3 individual molecules. Error bars are standard error of mean (S.E.M.).

### Chaperone mediated change on the average unfolding and refolding force of talin

To systematically investigated the mechanical stability of talin, we monitored the unfolding and refolding transition force of talin under varying force ramp protocols. Fig. 3A illustrates the mechanical stability of the talin in the presence and absence of p47, where the unfolding force of talin was measured by applying a constant pulling velocity of 9 pN/sec. The average unfolding force of talin in the absence of p47 was measured at 11.4±0.3 pN. However, in the presence of p47, this value increases significantly to 24.9 ± 1.68 pN, at a constant velocity of 9 pN/sec, indicating enhanced mechanical stability of talin (Fig. 3B). A similar trend in the unfolding force was observed at lower ramp rate of 1 pN/sec. Here we observed that the average unfolding force of talin was observed at 10.05±0.23 pN whereas it increases to 13.1±0.82 pN in the presence of p47, at a constant rate of 1 pN/sec, (Supp. Fig. 2). These results demonstrate the stabilizing effect of p47 on talin under mechanical stress. We further conducted a one-way ANOVA to assess the statistical significance of variations among the force data sets. The results demonstrated that the presence of p47 significantly influenced the average unfolding forces (Supp. Fig. 3).

**Figure 3:**
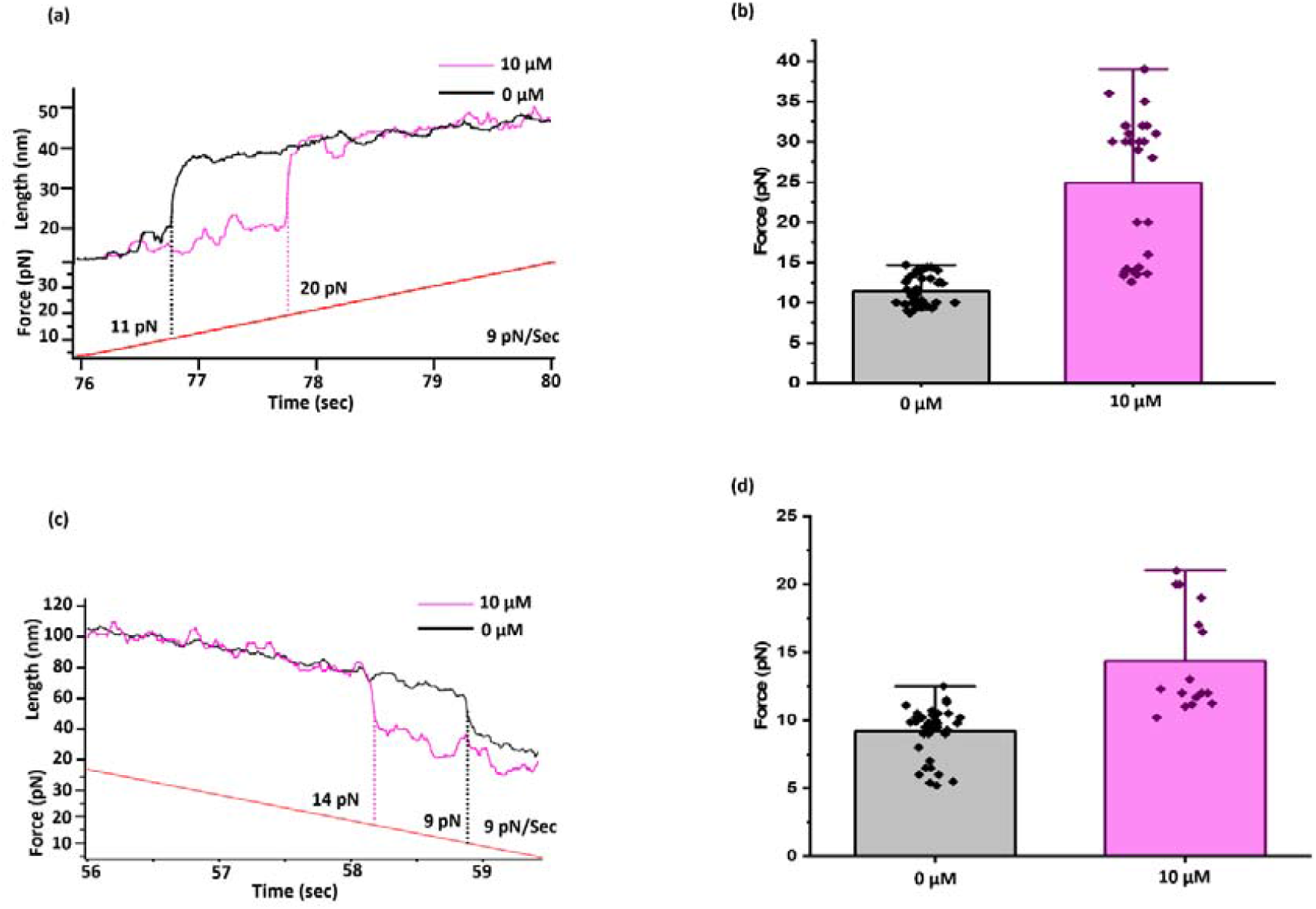
Average unfolding and refolding force of talin in the presence and absence of p47 at a constant pulling velocity of 9 pN/sec: **(A)** Representative unfolding traces of talin in the presence and absence of p47 at three constant different ramp rates of 9 pN/sec. **(B)** At a pulling velocity of 9 pN/sec, the average unfolding force of talin increased significantly from 11.4±0.3 pN to 24.9±1.68 pN , in the presence of p47 (pink). Each data points represents measurement of >3 individual molecules. Error bars are standard error of mean. **(C)** Representative traces of talin refolding in the presence of p47 were obtained at constant ramp rate of 9 pN/sec. **(D)** A comparable increase in the average refolding force was observed at a puling velocity of 9 pN/sec signifies the influence of p47 on the mechanical strength of talin. The average refolding force of talin increases from 9.12±0.3 to 14±0.94 pN, in the presence of p47 (pink) at a constant pulling velocity of 9 pN/sec. Each data points are taken by analysing >3 individual molecules. Error bars are standard error of mean.

Next, we examined the average refolding forces of talin at varying ramp rates in the presence of p47. Similar to its effect on unfolding behaviour, the refolding force of talin was significantly altered in the presence of p47 (Fig. 3C). At a ramp rate of 9 pN/sec, the refolding force increased from 9.12±0.3 to 14±0.94 pN in the presence of p47 (Fig. 3D). This increment was also consistent at another lower ramp rate of 1 pN/sec. Here we have found that the average refolding force of talin increased from 10.05±0.24 pN to 16.25±1.76 pN in the presence of p47, at a constant velocity of 1 pN/sec (Supp. Fig. 4). These findings further emphasize the stabilizing role of p47 on talin under mechanical stress. To validate the statistical significance of differences across various force data sets, we performed a one-way ANOVA. The analysis revealed that the changes in the average unfolding and refolding forces were statistically significant in the presence of p47 (Supp. fig. 5).

### The generation of extra mechanical work in the presence of p47

Next, to further explore the role of p47, we calculated the total mechanical work done associate with protein refolding under force. It is well known that the protein translocation at the edge of the molecular tunnel, such as the ER translocon or ribosomal translocon, generates mechanical work, and chaperones associated the tunnel can influence the substrate translocation under force^13,26^. The total amount of mechanical work output was determined by multiplying the step size of the protein with the applied refolding force^27,28^. The distribution curve showed that with increasing the force the mechanical work output is increasing, however the folding probability is decreasing at the higher force region. To account for this, we calculated the effective folding work done by multiplying the folding probability with the folding work. Remarkably, we found that in the absence of p47, talin required a maximum of 108.54 zJ of mechanical work to refold the protein at a particular force of 7 pN. However, in the presence of p47 it can generates maximum 199.75 zJ of mechanical work output at 12 pN (Fig. 4A). This additional mechanical work generated by p47 highlights its chaperone-like activity, which could play a crucial role in facilitating retrotranslocation of the polyprotein constructs from the ER lumen to the cytoplasm as proposed by our simple model (Fig. 4B). This model highlights the critical roles of p47 in retrotranslocation by boosting the mechanical work under force and provide critical support to translocate the substrate during retrotranslocation from ER lumen to the cytosol.

**Figure 4:**
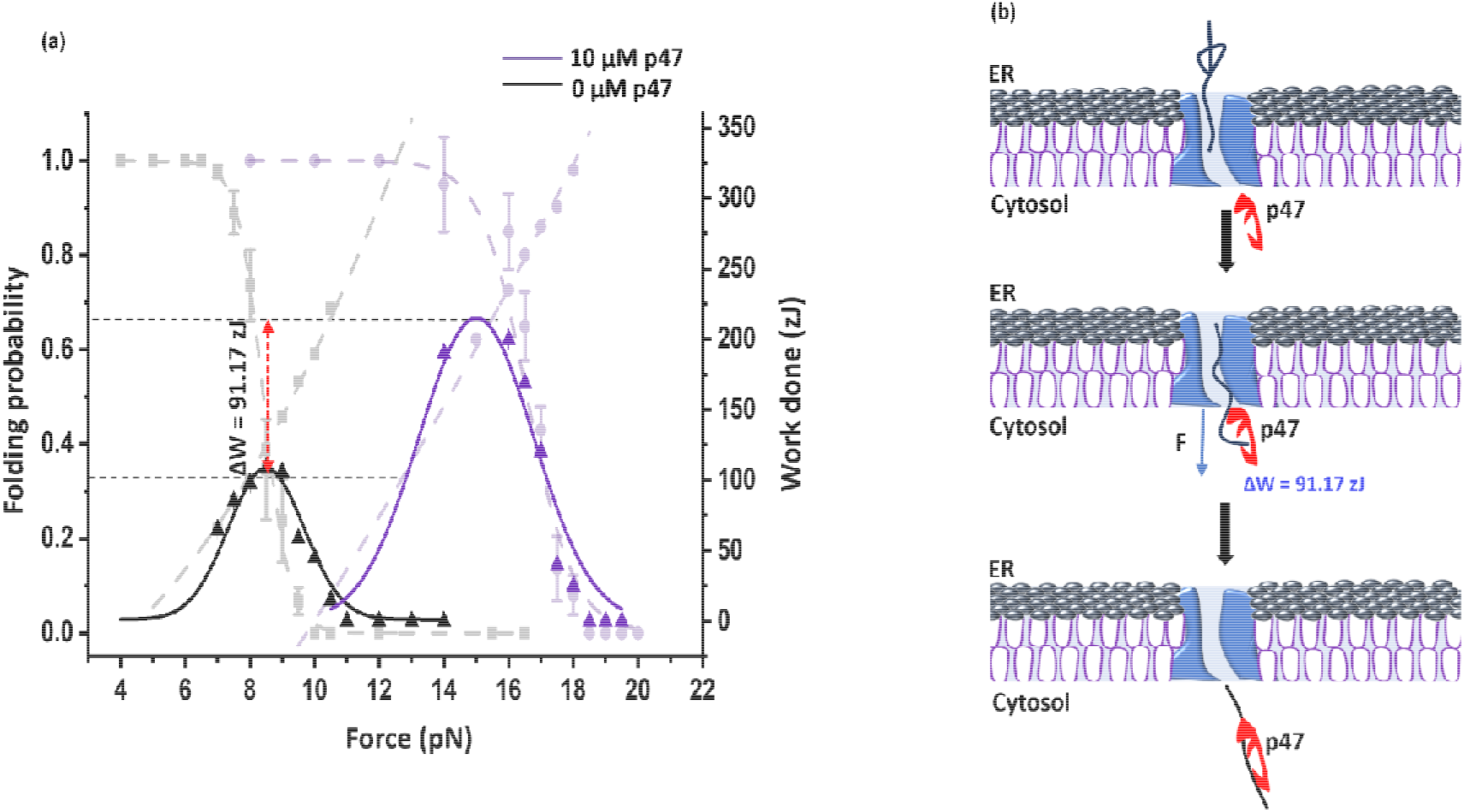
p47 mediated mechanical work output of talin and illustration of proposed model: **(A)** Total mechanical work done performed by talin was quantified as the product of folding probability with force and step size. In the absence of p47 (black) talin generated an optimal mechanical work done of 108.54 zJ whereas, its drastically increases up to 199.75 zJ in the presence of p47(purple). Error bars are standard error of mean (S.E.M.). **(B)** This figure depicts the engagement of p47 in the extraction of proteins from the ER lumen to the cytoplasm under mechanical force. Here p47 acta as a chaperone like property and boost mechanical work output, facilitating the polypeptide to translocate through the pore.

## Discussion

ER-associated degradation (ERAD) is a crucial quality control pathway that eliminates misfolded or aberrant proteins from the endoplasmic reticulum (ER) to maintain cellular proteostasis^29–31^. A central step in this process, retrotranslocation, extracts proteins from the ER lumen to the cytosol, where they are targeted for proteasomal degradation, ensuring the fidelity of the secretory pathway^32^. Despite its significance, the mechanism driving the retrotranslocation of the proteins and the driving force that mediates the substrate extraction remain incompletely understood. Using single molecule force spectroscopic technique, we have tried to understand the roles of a cofactor p47, hypothesized to act as a chaperone, in facilitating the retrotranslocation of the unfolded polypeptides.

Our results demonstrates that p47 significantly enhances the mechanical stability of the substrate talin and shifting its half-point force to higher values and thereby increasing its mechanical resilience. This shift indicates that p47 promotes the ability of talin to refold at higher applied forces, suggesting a protective role for the cofactor under mechanical stress. Using force ramp protocols, we demonstrated that p47 significantly enhances that mechanical strength of talin. At a loading rate of 9 pN/sec, the unfolding force increased from 11.4 ± 0.3 pN to 24.9 ± 1.68 pN, while the refolding force also increase from 9.12 ± 0.3 pN to 14 ± 0.94 pN (Fig. 3B and 3D). These findings underscore the critical role of p47 in stabilizing talin under mechanical stress, highlighting its ability to modulate the mechanical stability of talin.

In this context, force is recognized as a critical factor in protein translocation through molecular channels. Previous studies have demonstrated that mechanical force can play dual role by guiding the protein through the channel and preventing backsliding, as previously discussed by the Brownian ratchet mechanism^33–37^. Mechanical forces generated by chaperones like Hsp70, or nascent chain folding play a crucial role in regulating ribosome elongation and protein translocation by alleviating stalling and ensuring efficient passage through the translocon pore^38–40^. Notably, these force can generate strong entropic pulling forces (∼ 46 pN) to extract trapped polypeptides, highlighting a universal mechanism for regulating protein translocation through the translocon pore^41^. Interestingly, these forces appear to work synergistically with protein folding dynamics to optimize mechanical work during extraction. In this context, it still unclear whether such force transmission could occur during retrotranslocation through the tunnel to facilitate the protein translocation. However, it is plausible that certain chaperones or cofactors play crucial role by generating extra pulling forces to extract the proteins from the tunnel that could aid in directing the protein toward cytosolic degradation or refolding pathways, might provide vital rescue mechanism to ensure proteostasis. Here, our comprehensive analysis showed that p47 enhances the maximum mechanical work output of ∼199.75 zJ under mechanical stress, underscoring its efficiency to extract the polypeptide through the tunnel from ER to cytosol. These findings suggest a model where p47 associates with the ER translocon and directly engages with the polypeptide chain and regulates translocation through a molecular pore by generating additional mechanical force and work out put. In summary, our study reveals a novel chaperone-like activity of p47, facilitating substrate stabilization and mechanical extraction during retrotranslocation, thereby offering new mechanistic insights into protein quality control processes.

## Analysis

All the data analysis and acquisition and analysis were performed with Origin pro and Igor Pro 8.0 software (Wavemeters) software.

## Author contribution

S.H. and D.C. designed the project. D.C. performed the experiment, analysed the data and wrote the manuscript.

## Acknowledgements

We thank the department of Chemistry, Ashoka University, S.N.Bose National Centre for Basic Science and Technical Research Centre of S.N.Bose National Centre for Basic Science for the support and funding.

